# Anti-tubercular activity of novel 4-anilinoquinolines and 4-anilinoquinazolines

**DOI:** 10.1101/570002

**Authors:** Christopher R. M. Asquith, Neil Fleck, Chad D. Torrice, Daniel J. Crona, Christoph Grundner, William J. Zuercher

**Author notes:** Corresponding authors: (C. Grundner), (W. J. Zuercher).

## Abstract

We screened a series of 4-anilinoquinolines and 4-anilinoquinazolines and identified novel inhibitors of *Mycobacterium tuberculosis* (*Mtb*). The focused 4-anilinoquinoline/quinazoline scaffold arrays yielded compounds with high potency and the identification of 6,7-dimethoxy-*N*-(4-((4-methylbenzyl)oxy)phenyl)quinolin-4-amine (**34**) with an MIC_90_ value of 0.63-1.25 μM. We also defined a series of key structural features, including the benzyloxy aniline and the 6,7-dimethoxy quinoline ring, that are important for *Mtb* inhibition. Importantly the compounds showed very limited toxicity and scope for further improvement by iterative medicinal chemistry.

---

*Mycobacterium tuberculosis* (*Mtb*), the causative agent of tuberculosis (TB) in humans,^1^ infects nearly a third of the earth’s population and caused 1.6 million worldwide deaths in 2017.^2^ With nearly ten million new cases of active disease each year, TB is now the leading cause of death from infectious disease globally.^2^ Current therapeutic strategies involve the use of a combination of anti-microbial agents including ethambutol, isoniazid, pyrazinamide and rifampicin (Fig. 1).^3^ However, more than 5% of *Mtb* infections now involve multidrug-resistant (MDR-TB) and extensively drug-resistant (XDR-TB) *Mtb* strains. MDR-TB is associated with a 50% mortality rate whereas XDR-TB is nearly always fatal.^4^ There is an urgent need for new therapeutic strategies.

**Figure 1.**
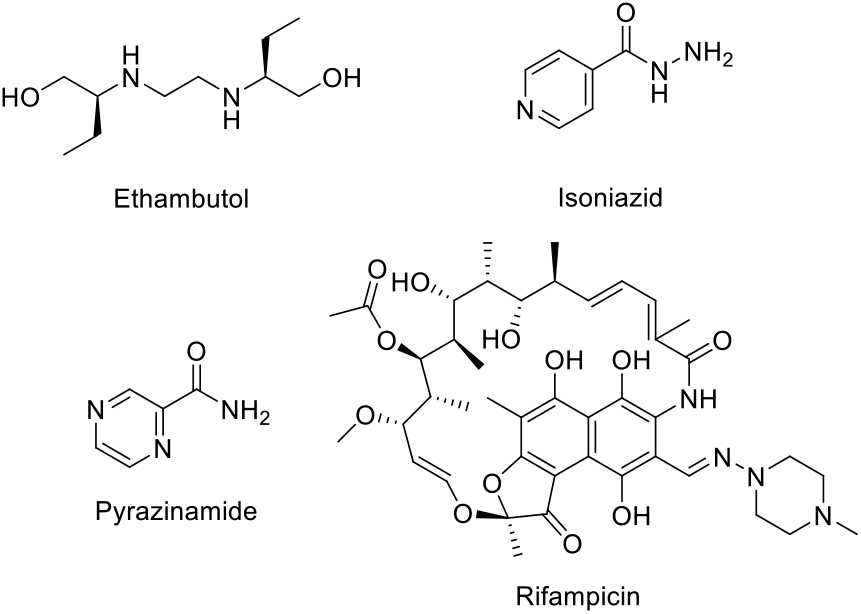
Current therapeutic strategies for treatment of *Mycobacterium tuberculosis* infections.

Human protein kinases are pharmacologically tractable enzymes targeted by more than three dozen approved medicines.^5^ Hundreds of additional kinase inhibitors are under clinical and preclinical investigation. There is growing recognition that pathogen kinases may be targeted in the treatment of infectious diseases.^6–7^ Considering the conserved ATP-binding site across species, we looked to screen collections of ATP-competitive inhibitors of human kinases for their anti-tubercular activity.

To identify new chemical starting points against *Mtb* we looked to lapatinib, gefitinib, and erlotinib as starting points which have recently been revealed to have activity against *Mtb* (Fig. 2.).^5,8^ We tested the activity of lapatinib, gefitinib, and erlotinib against *Mtb* by measuring luminescence and growth on solid medium across a series of four two-fold dilutions starting at 20 μM (Tab 1).^9–10^ The reduction of visible growth on solid medium demonstrated the compounds to be bactericidal.

**Figure 2.**
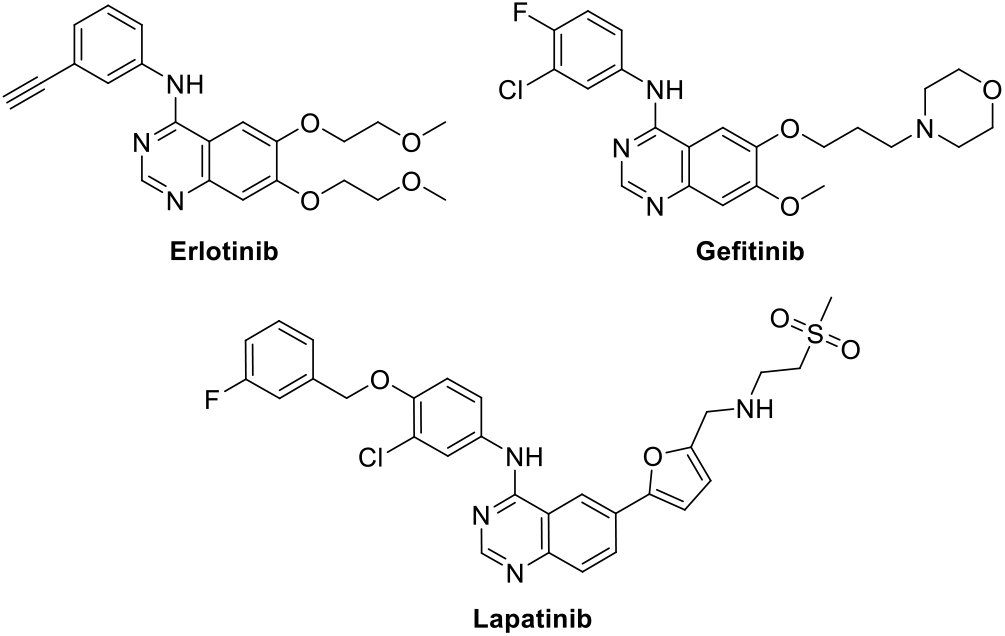
Structures of clinical quinazolines.

**Table 1.**
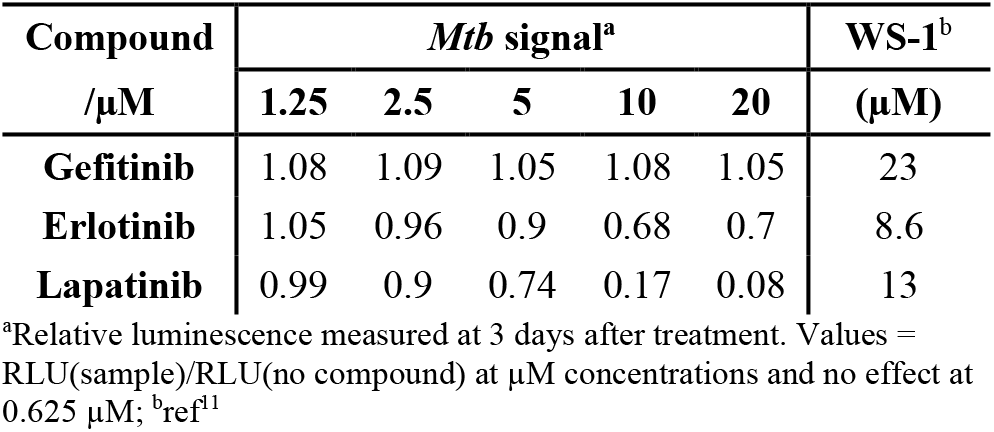
Results of clinical inhibitors.

| Compound<br>/$\mu\text{M}$ | *Mtb* signal<sup>a</sup> | | | | | WS-1<sup>b</sup><br>($\mu\text{M}$) |
| --- | --- | --- | --- | --- | --- | --- |
|  | 1.25 | 2.5 | 5 | 10 | 20 |  |
| **Gefitinib** | 1.08 | 1.09 | 1.05 | 1.08 | 1.05 | 23 |
| **Erlotinib** | 1.05 | 0.96 | 0.9 | 0.68 | 0.7 | 8.6 |
| **Lapatinib** | 0.99 | 0.9 | 0.74 | 0.17 | 0.08 | 13 |

<sup>a</sup>Relative luminescence measured at 3 days after treatment. Values = RLU(sample)/RLU(no compound) at $\mu\text{M}$ concentrations and no effect at 0.625 $\mu\text{M}$ ; <sup>b</sup>ref<sup>11</sup>

Gefitinib treatment had no effect relative to the absence of compound. Erlotinib induced a modest effect that appeared to plateau at a signal of approximately 70 %. In contrast, lapatinib showed activity even at 5 μM and reduced the relative *Mtb* signal to below 10 % at 20 μM. This result suggested that the 4-benzyloxy aniline substituent might be important for anti-Mtb activity. These compounds demonstrate only limited toxicity in a human skin fibroblast cell line (WS-1) counter screen.^11^

To further explore quinazoline *Mtb* activity, we profiled several focused arrays of compounds to probe the structure activity relationships of the quinoline/quinazoline. We hence synthesized a series of compounds (**1-34**) following up on the results listed in table 1, exploring the 4-anilinoquinoline and 4-anilinoquinazoline scaffolds through nucleophilic aromatic displacement of 4-chloroquin(az)olines. (Sch. 1) We were able to access products in good to excellent yields (55-91 %) consistent with previous reports and without protection of the alcohol substituted quin(az)oline starting material.^11–13^

**Scheme 1.**
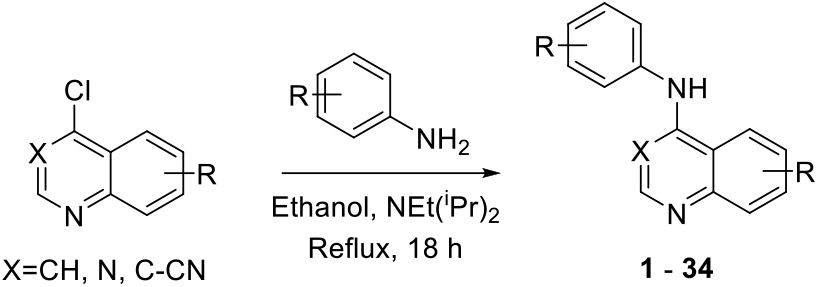
General synthetic procedure

The first set of compounds probed a replacement of the 6-postion morpholine segment of gefitinib with a simple alcohol on the lapatinib scaffold (Tab. 2).^20^ Although neither the 6-OH (**1**) or 7-OH (**2**) quinazoline showed appreciable activity, the 6,7-dihydroxy compound (**3**) began to inhibit *Mtb* growth at higher concentrations. The analogous set of methoxy-substituted compounds (**4-6**) had very similar activity profiles. The 6-OH quinoline (**7**) showed improved activity relative to the matched quinazoline (**1**). The inclusion of fluorine substitution on the phenyl ring distal to the quinazoline (**8-10**) led to markedly increased activity for all three isomers. At 20 μM, **8-10** all reduced the relative *Mtb* signal to 25-37 %, and a modest but discernable reduction in signal was observed at 1.25 μM. Interestingly, the effect of fluorine substitution led to a different activity pattern from changing quinazoline ring substitution as the modification of **1** to the 7-OH (**11**) or 6,7-(OH)_2_ (**12**) resulted in a significant loss of activity, even at 20 μM. These results demonstrated that *Mtb* activity was sensitive to changes at multiple parts of the template and that these changes were not necessarily additive. We counter screened **1-12** in human skin fibroblast cells (WS-1) and observed very limited toxicity with **3** and **7** the only compounds in the single digit micromolar range (IC_50_ = 8.8 and 2.5 μM respectively).^15^

**Table 2.** Results of alcohol replacement of lapatinib and gefitinib (**1-12**).

| Compound | X | R<sup>1</sup> | R<sup>2</sup> | R<sup>3</sup> | *Mtb* signal<sup>a</sup> ($\mu\text{M}$) | | | | | WS-1<sup>b</sup><br>($\mu\text{M}$) |
| --- | --- | --- | --- | --- | --- | --- | --- | --- | --- | --- |
|  |  |  |  |  | 1.25 | 2.5 | 5 | 10 | 20 |  |
| **1** | N | OH | H | H | 0.95 | 0.94 | 0.96 | 0.90 | 0.80 | 26 |
| **2** | N | H | OH | H | 0.90 | 0.90 | 0.89 | 0.88 | 0.89 | >100 |
| **3** | N | OH | OH | H | 0.95 | 0.93 | 0.91 | 0.76 | 0.69 | 8.8 |
| **4** | N | OMe | H | H | 1.04 | 1.07 | 0.89 | 0.83 | 0.62 | >100 |
| **5** | N | H | OMe | H | 1.05 | 1.12 | 0.99 | 0.87 | 0.64 | >100 |
| **6** | N | OMe | OMe | H | 1.04 | 1.04 | 0.83 | 0.77 | 0.6 | >100 |
| **7** | CH | OH | H | H | 0.95 | 0.86 | 0.66 | 0.42 | 0.37 | 2.5 |
| **8** | N | OH | H | 4-F | 0.86 | 0.76 | 0.64 | 0.44 | 0.32 | >100 |
| **9** | N | OH | H | 3-F | 0.76 | 0.64 | 0.49 | 0.39 | 0.25 | 12 |
| **10** | N | OH | H | 2-F | 0.89 | 0.82 | 0.63 | 0.46 | 0.28 | 28 |
| **11** | N | H | OH | 4-F | 0.94 | 1.01 | 0.90 | 0.89 | 0.83 | >100 |
| **12** | N | OH | OH | 4-F | 1.04 | 1.14 | 1.00 | 0.89 | 0.76 | >100 |

<sup>a</sup>Relative luminescence measured at 3 days after treatment. Values = RLU(sample)/RLU(no compound) at $\mu\text{M}$ concentrations and no effect at 0.625 $\mu\text{M}$ ; <sup>b</sup>IC<sub>50</sub> (mean average n=4), 48 h

The next set of quin(az)olines profiled was prepared to explore the contributions from both the aniline and the core heterocycle (Tab. 3).^11,16^ The larger 3,4,5-trimethoxyphenyl aniline was employed on several diversely substituted quinolines and quinazolines (**13-18**) which led to no observable activity except when paired with the 6,7-(OCH_2_CH_2_OMe)_2_-quinazoline of erlotinib (**19**) which had only slight activity at 20 μM. On the other hand, incorporation of the aniline fragment from erlotinib (3-ethynylphenyl) did yield several active compounds, including the quinoline analog of erlotinib (**20**). The erlotinib aniline with a 6,7-(OMe)_2_-substituted quinoline core (**21**) or quinazoline (**22**) showed activity but not with the 3-cyanoquinoline (**23**). The analogous 3-bromophenyl aniline compounds (**24-26**) showed higher activity than the paired 3-ethynylphenyl compounds. Finally, the aniline substitution from lapatinib (3-Cl-4-(2-F-PhO)Ph) yielded a marginally active quinazoline (**27**) and a highly active quinoline (**28**) that had 10 % *Mtb* signal at 20 μM. We counter screened **13-28** in WS-1 cells and observed limited toxicity in most compounds. However, the bromine substitution appeared to increase toxicity (**24-26**) along with the lapatinib derivatives (**27-28**).

**Table 3.** Matched pair comparison of structures similar to erlotinib and lapatinib (**13-28**).

| Compound | R<sup>1</sup> | R<sup>2</sup> | X | R<sup>3</sup> | *Mtb* signal<sup>a,b</sup> | | | WS-1<sup>c</sup><br>($\mu\text{M}$) |
| --- | --- | --- | --- | --- | --- | --- | --- | --- |
| | | | | | 5 $\mu\text{M}$ | 10 $\mu\text{M}$ | 20 $\mu\text{M}$ | |
| **13** | CF<sub>3</sub> | H | CH | 3,4,5-(OMe)<sub>3</sub> | 1.23 | 1.14 | 1.21 | >100 |
| **14** | CF<sub>3</sub> | H | N | 3,4,5-(OMe)<sub>3</sub> | 1.25 | 1.12 | 1.06 | >100 |
| **15** | Br | H | CH | 3,4,5-(OMe)<sub>3</sub> | 1.35 | 1.41 | 1.28 | >100 |
| 16 | OMe | OMe | CH | 3,4,5-(OMe)<sub>3</sub> | 1.26 | 1.11 | 1.06 | >100 |
| 17 | OMe | OMe | N | 3,4,5-(OMe)<sub>3</sub> | 1.03 | 1.01 | 0.94 | >100 |
| 18 | OMe | OMe | CN | 3,4,5-(OMe)<sub>3</sub> | 1.03 | 1.01 | 0.97 | >100 |
| 19 | 6,7-(OCH<sub>2</sub>CH<sub>2</sub>OMe)<sub>2</sub> |  | N | 3,4,5-(OMe)<sub>3</sub> | 1.02 | 0.96 | 0.87 | >100 |
| 20 | 6,7-(OCH<sub>2</sub>CH<sub>2</sub>OMe)<sub>2</sub> |  | CH | 3-Ethynyl | 0.88 | 0.79 | 0.58 | >100 |
| 21 | OMe | OMe | CH | 3-Ethynyl | 1.02 | 0.89 | 0.67 | >100 |
| 22 | OMe | OMe | N | 3-Ethynyl | 0.93 | 0.83 | 0.69 | >100 |
| 23 | OMe | OMe | CN | 3-Ethynyl | 1.08 | 1.03 | 0.98 | >100 |
| 24 | OMe | OMe | CH | 3-Bromo | 1.03 | 0.90 | 0.6 | 9.6 |
| 25 | OMe | OMe | N | 3-Bromo | 0.98 | 0.71 | 0.65 | 3.9 |
| 26 | OMe | OMe | CN | 3-Bromo | 0.88 | 0.56 | 0.37 | 11 |
| 27 | OMe | OMe | N | 3-Cl-4-(2-F-PhO) | 0.92 | 0.74 | 0.80 | 1.1 |
| 28 | OMe | OMe | CH | 3-Cl-4-(2-F-PhO) | 0.78 | 0.22 | 0.10 | 11 |

<sup>a</sup>Relative luminescence was measured at 3 days after treatment. Values = RLU(sample)/RLU(no compound); <sup>b</sup>None of the compounds reduced the relative *Mtb* signal below 95 % at 1.3 or 2.5 μM; <sup>c</sup>IC<sub>50</sub> (mean average n = 4), 48 h

A subsequent set of compounds explored features of the 4-benzyloxyaniline portion of the quinoline template (Tab. 4).^17–18^ Variation of the ether linkage to an amide and addition of a hydroxy on the aniline portion revealed an activity pattern where 6,7-dimethoxy substitution with the benzylic arm was active (**29-30**). In contrast, truncation of the benzyl also removed the activity as in acetamide **31**, which had no effect on *Mtb*. The 6-methoxy (**32**) and 7-methoxy (**33**) showed no reduction in *Mtb* signal at 20 μM. However, switching back to the 4-methyl benzyl ether linked compound **34** yielded the most potent activity observed with any compound in the present study, with a robust signal observed even at 1.25 μM. The *Mtb* MIC_90_ for **34** was in the 0.63-1.25 μM range. However, the kill curve plateaued at 5 μM, and no improved killing was observed at higher doses (98 % inhibition at 5 μM; 98 % inhibition at 20 μM) (Fig. 3).

**Table 4.** Matched pair comparison of benzyloxyaniline.

| Compound | R<sup>1</sup> | R<sup>2</sup> | R<sup>3</sup> | R<sup>4</sup> | *Mtb* signal<sup>a</sup> (μM) | | | | | WS-1<sup>b</sup> (μM) |
| --- | --- | --- | --- | --- | --- | --- | --- | --- | --- | --- |
|  |  |  |  |  | 1.25 | 2.5 | 5 | 10 | 20 |  |
| 29 | OMe | OMe | OH | A | 1.14 | 1.09 | 1.13 | 1.05 | 0.45 | 16 |
| **30** | OMe | OMe | OH | **B** | 1.10 | 1.12 | 1.06 | 0.72 | 0.25 | 2.6 |
| **31** | OMe | OMe | H | **C** | 1.21 | 1.10 | 1.11 | 1.07 | 1.02 | 9.7 |
| **32** | OMe | H | OH | **A** | 1.11 | 1.17 | 1.14 | 1.28 | 0.95 | 10 |
| **33** | H | OMe | OH | **A** | 1.13 | 1.11 | 1.09 | 1.11 | 0.96 | 4.7 |
| **34** | OMe | OMe | H | **D** | 0.43 | 0.17 | 0.10 | 0.08 | 0.09 | 5.4 |

<sup>a</sup>Relative luminescence was measured at 3 days after treatment. Values = RLU(sample)/RLU(no compound); <sup>b</sup>IC<sub>50</sub> (mean average n=4), 48 h

**Figure 3.**
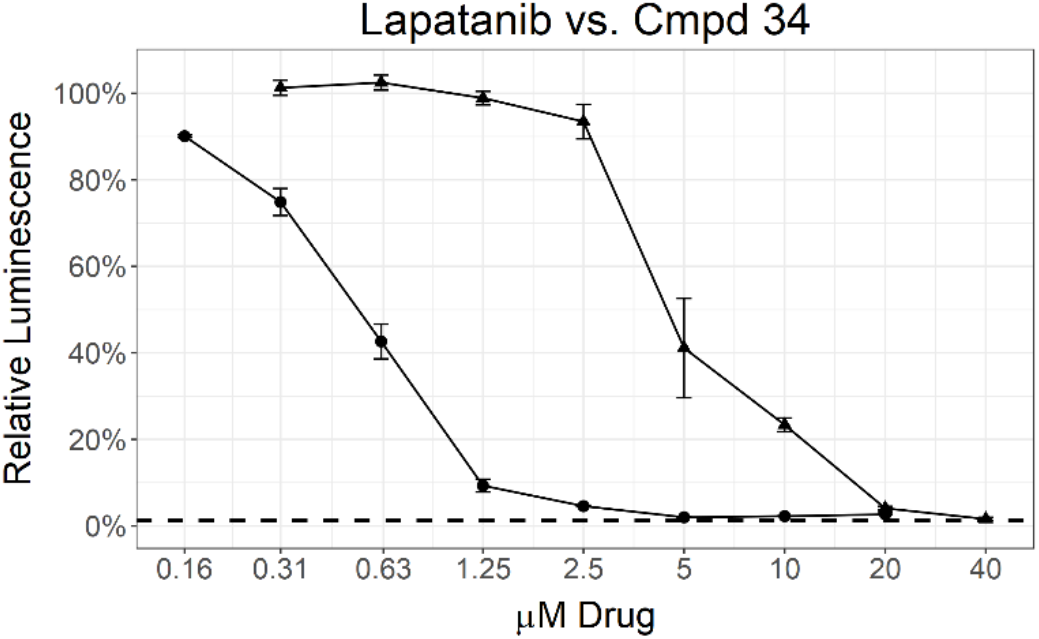
MIC determination by two different assays for **34** (circle) and lapatanib (triangle). Data points represent the mean of 3 biological replicates with standard deviation. The dashed line labeled 1% inoculum represents an inoculation of 1 % of the number of cells used for compound testing. This control was used to determine 99 % inhibition of growth.

As with the other compound sets, we evaluated **29-34** in human skin fibroblast cells (WS-1) and observed moderate toxicity in the single digit micromolar range for most compounds.^20^ Importantly, the anti-*Mtb* effects of compounds appeared to be divergent from the toxic effects in WS-1 cells, suggesting that the *Mtb* effects were not driven by nonspecific cytotoxicity. The most potent anti-*Mtb* compound 34 had WS-1 IC_50_ = 5.4 μM, substantially higher than its *Mtb* MIC_90_ value and within threefold of the IC_50_ values for erlotinib and lapatinib. This result demonstrated that, in the human WS-1 cell line, **34** behaved comparably to two approved medicines.

These structure activity relationships between *Mtb* activity and the 4-anilinoquinoline/quinazoline scaffold have the potential to inform a medicinal chemistry strategy for enhanced *Mtb* activity. The most sensitive structural changes were found to be in the ring appended to the aniline rather than in the quin(az)oline core.

This body of work provides a number of exciting starting points for further optimization, with limited non-specific toxicity. However, the failure to achieve complete parasite kill led us to deprioritize the series due to the potential for resistance to develop. The mechanism of anti-*Mtb* activity of the quin(az)olines has yet to be defined. These compounds were originally prepared as inhibitors of human kinases targeting the ATP-binding site, a rational starting hypothesis for the mechanism of action is that the effects of these compounds are mediated by inhibition of *Mtb* kinases. However, it is possible that the observed phenotypes may originate from modulation of other, non-kinase ATP-binding proteins in the organism.

Gefitinib, erlotinib, and lapatinib have previously been reported to inhibit the intracellular growth of *Mtb*. Multiple lines of evidence were described suggesting inhibition of the host target epidermal growth factor receptor (EGFR) was responsible for this activity.^15^ However our results demonstrate that proteins within the pathogen itself may be targeted as well. The benzyl substituent present in the molecule showed a pivotal effect to potency, as is highlighted by the enhanced activity of **34** relative to **30**. The present results help define a de-risked medicinal chemistry trajectory towards anti-tubercular compounds with targets in both the host and the parasite itself. Such dual acting compounds might offer advantages in efficacy and/or reduction in propensity for resistance.

## Acknowledgments

The SGC is a registered charity (number 1097737) that receives funds from AbbVie, Bayer Pharma AG, Boehringer Ingelheim, Canada Foundation for Innovation, Eshelman Institute for Innovation, Genome Canada, Innovative Medi-cines Initiative (EU/EFPIA) [ULTRA-DD grant no. 115766], Janssen, Merck KGaA Darmstadt Germany, MSD, Novartis Pharma AG, Ontario Ministry of Economic Development and Innovation, Pfizer, São Paulo Research Foundation-FAPESP, Takeda, and Wellcome [106169/ZZ14/Z]. C. G. is supported by R01 AI117023 from the NIH/NIAID. We are grateful Dr. Brandie Ehrmann for LC-MS/HRMS support provided by the Mass Spectrometry Core Laboratory at the University of North Carolina at Chapel Hill.

